# Improved characterisation of clinical text through ontology-based vocabulary expansion

**DOI:** 10.1101/2020.07.10.197541

**Authors:** Luke T Slater, William Bradlow, Simon Ball, Robert Hoehndorf, Georgios V Gkoutos

## Abstract

**Background:** Biomedical ontologies contain a wealth of metadata that constitutes a fundamental infrastructural resource for text mining. For several reasons, redundancies exist in the ontology ecosystem, which lead to the same concepts being described by several terms in the same or similar contexts across several ontologies. While these terms describe the same concepts, they contain different sets of complementary metadata. Linking these definitions to make use of their combined metadata could lead to improved performance in ontology-based information retrieval, extraction, and analysis tasks.

**Results:** We develop and present an algorithm that expands the set of labels associated with an ontology class using a combination of strict lexical matching and cross-ontology reasoner-enabled equivalency queries. Across all disease terms in the Disease Ontology, the approach found **51**,**362** additional labels, more than tripling the number defined by the ontology itself. Manual validation by a clinical expert on a random sampling of expanded synonyms over the Human Phenotype Ontology yielded a precision of **0.912**. Furthermore, we found that annotating patient visits in MIMIC-III with an extended set of Disease Ontology labels led to semantic similarity score derived from those labels being a significantly better predictor of matching first diagnosis, with a mean average precision of **0.88** for the unexpanded set of annotations, and **0.913** for the expanded set.

**Conclusions:** Inter-ontology synonym expansion can lead to a vast increase in the scale of vocabulary available for text mining applications. While the accuracy of the extended vocabulary is not perfect, it nevertheless led to a significantly improved ontology-based characterisation of patients from text in one setting. Furthermore, where run-on error is not acceptable, the technique can be used to provide candidate synonyms which can be checked by a domain expert.

## Background

Metadata are a fundamental feature of biomedical ontologies, describing a wealth of natural language information in the form of labels and descriptions [1]. OWL ontologies implement metadata in the form of annotation properties, and these can be used to describe multiple natural language labels for a single term. Open Biomedical Ontologies (OBO) [2] define a series of conventional annotation properties that can be used for the expression of labels and synonyms.

These features are widely used; as of 2017 the Human Phenotype Ontology (HP) [3] contained 14,328 synonyms for 11,813 classes [4]. Because such labels are associated with ontology terms, ontologies constitute a controlled domain vocabulary.

The role of a controlled domain vocabulary makes ontologies a valuable resource for text mining, particularly in information retrieval and extraction tasks [5]. Furthermore, association of entities described in text with ontologies enables their integration with other datasets annotated by those ontologies, as well as caters the application of ontology-based analysis techniques such as semantic similarity, semantic rule-mining, or relational machine learning.

However, due to limitations of resources for expert curation of ontologies and the sheer scale of their contents, the labels obtainable from single ontologies are not exhaustive. Combined with the tendency for alternative presentation of semantically equivalent concepts in biomedical text [6], ontology labels are not always a good fit for text corpora that discuss the same concepts [7]. By expanding the set of synonyms in an ontology, particularly with synonyms that provide a better fit for text corpora, the performance of natural language processing tasks that depend on them can be improved.

This potential is reflected by previous work in the field. One approach that used analysis of existing synonyms across ontology hierarchy to determine new synonyms reported an increase in performance of a task retrieving articles from a literature repository [8]. Another rule-based synonym expansion approach to extending the Gene Ontology showed improved performance in named entity recognition (NER) tasks [9]. A combined machine-learning and rule-based approach to learning new HP synonyms from manually annotated PubMed abstracts improved performance of an annotation task over a gold standard text corpus [10].

Ontology-based annotation software such as OBO Annotator [11], ConceptMapper [12], and the NCBO Annotator [13] contain routines to consider rule-based morphological and positional transformations of terms to increase NER recall. Parameters that control the use of these features have a strong influence on annotation performance [14]. Previous work has also investigated synonym acquisition and derivation for the purposes of improving the performance of lexical ontology matching and alignment tasks [15]. Outside of automated synonym generation, organised efforts have been made to manually extend an ontology’s synonyms for a particular purpose. For example, HP was expanded with layperson synonyms to enable its use in applications that interact directly with patients [16].

However, no work to our knowledge has considered linking different ontology terms for the purposes of vocabulary expansion. Many biomedical concepts are described by the same terms in equivalent or similar contexts across several ontologies. For example, terms describing hypertension exist in many ontologies and medical terminologies. The *hypertension* (HP:0000822) term describes the condition in the context of a phenotype, while *hypertension* (DOID:10763) from the Disease Ontology (DO) [17] describes it in the context of a disease. Specific-disease or application ontologies also extend upon definitions provided by general domain ontologies. For example, the Hypertension Ontology (HTN) [18] extends the HP and DO hypertension classes, adding additional information including labels. Furthermore, the subtle distinctions between concepts that biomedical ontologies capture, including phenotype versus disease, do not necessarily influence many of the commonly applied text mining tasks, because these contexts share the same labels.

We hypothesise that because ontologies are constructed with different loci, ontologies that define terms describing the same concepts will contain different, but valid, synonyms for a particular context. These loci consist in contexts, domain experts, and source material. By considering all of these terms, we can construct extended vocabularies that may improve the power of ontology-based text mining tasks. In this paper, we describe and implement a synonym expansion approach that combines lexical matching and semantic equivalency to obtain new synonyms for biomedical concepts. We use the approach to extend several ontology vocabularies, and evaluate them both manually, and in an ontology-based patient characterisation task.

## Results

The synonym expansion algorithm is available as part of the Komenti text mining framework, which is available under an open source licence at https://github.com/reality/komenti, while the files used for validation are available at https://github.com/reality/synonym_expansion_validation.

**Algorithm**

We developed a synonym expansion algorithm that derives additional synonyms for a class by matching it with classes from other ontologies, making use of the AberOWL ontology reasoning framework [19]. The algorithm performs the following process, for each class provided as input (in this context, ‘every ontology’ is any of the ontologies that are included in AberOWL):

1. Extract the labels and synonyms of any classes in any ontology with a label or synonym that exactly matches the first label of the input class.
2. Run an equivalency query against every ontology using the Internationalised Resource Identifier (IRI) of the input class, extracting labels and synonyms for any classes returned.
3. Of the candidate synonyms produced by the first two steps, discard any that were:
  - Defined in ontologies that were found to produce incorrect synonyms.
  - Have the form of a term identifier.
  - Contain the input class label as a substring.

The algorithm uses two different methods for identifying matching classes. Strict lexical matching is used to identify otherwise unlinked terms that contain a label which is the same as the first label of the input class. Mapping terms across ontologies via shared labels or metadata is a well established technique used in ontology alignment [20]. Cross-ontology equivalency queries are used to obtain additional classes in the case that ontologies describe semantically equivalent classes, but do not share the same label. This can occur due to inferred semantic equivalency, ontologies becoming out of sync, or omission of annotation properties in a referencing class. Only the first label for the input class is used, because additional labels and synonyms were found to less uniquely identify the class in question, and lead to more incorrect candidate synonyms.

After the main matching stage, the set of labels is pruned down to remove incorrect values. Some ontologies include term identifiers as labels which cannot be exploited by text-mining applications. Therefore, candidate synonyms that contained a colon or underscore were removed. The algorithm also removes labels sourced from GO-PLUS [21], MONDO [22], CCONT [23], and phenX [24], because we found these ontologies consistently produced incorrect synonyms. We also removed labels that include the input label as a substring, as these add no value to NER systems (as the smaller string would match, making the longer string redundant).

### Ontology Expansion

We applied the vocabulary expansion algorithm to all 9,908 subclasses of *disease* (DOID:4) in the Disease Ontology (DO). DO itself asserts 24,878 labels and synonyms for these classes. The expanded DO vocabulary contained 76,240 labels and synonyms. We also applied the algorithm to the 14,406 non-obsolete subclasses of *Phenotypic abnormality* (HP:0000118) in HP. HP itself asserts 29,805 labels and synonyms. The number of labels and synonyms following expansion was 54,765. Therefore, the algorithm found 24,960 additional synonyms across HP.

For the DO term *hypertension* (DOID:10763), 28 labels and synonyms were found. 3 of these were from DO itself. The algorithm found 70 synonyms not including the word ‘hypertension’. Of these, 56 were obtained via lexical matching, and 14 by equivalency query. The sources of these synonyms is summarised in Table 1. After making the list unique, there were 28 labels and synonyms. Therefore, the algorithm found 25 new synonyms.

**Table 1.** Source of the 70 non-unique synonyms found for the term *hypertension* (DOID:10763) per-ontology. Of these synonyms, 28 were unique. Bracketed numbers, where given, are the labels found by the equivalency method only (in these cases, lexical matches were made for multiple classes in the ontology).

| Ontology | Source | Number of Synonyms |
| --- | --- | --- |
| GWAS_EFO_SKOS | Lexical | 16 |
| MESH | Lexical | 4 |
| HTN [18] | Lexical, Equivalency | 3 (2) |
| CRISP | Lexical | 1 |
| CCTOO [25] | Lexical | 6 |
| ONTONEO | Lexical, Equivalency | 4 (3) |
| NCIT [26] | Lexical | 6 |
| COSTART | Lexical | 7 |
| BAO | Lexical, Equivalency | 2 |
| CSSO | Lexical | 2 |
| ODAE | Lexical, Equivalency | 2 |
| DO | — | 3 |
| DTO | Equivalency | 2 |

In this example, there were no synonyms uniquely found via equivalency, however, if we use *bradycardia* as the input class, we can identify two new synonyms from PhenomeNET [27], *bradyrhythmia* and *reduced heart rate*, which were not otherwise obtained via lexical matching. This is because PhenomeNET establishes a semantic equivalency between *decreased heart rate* (MP:0005333), which does not share its first label with the HP class.

### Manual Validation

To evaluate the correctness of synonyms in the expansion of HP, a clinical expert manually validated the correctness of 866 synonyms found for 500 randomly selected terms. Table 2 summarises the results, which show a precision of 0.912. 195 terms were marked as ambiguous, in the case that the synonyms were in a foreign language or the clinician did not have enough expertise of the term to determine whether the synonym was correct.

**Table 2.** Metrics for clinical expert validation of 866 generated synonyms for 500 terms. Synonyms already included in HP were not included in the validation. Synonyms were marked ambiguous if not English, or if the validator did not have enough expertise to confidently judge it.

| Terms | Total Synonyms | TP | FP | Ambiguous | Precision |
| --- | --- | --- | --- | --- | --- |
| 500 | 866 | 613 | 59 | 195 | 0.912 |

### Information Retrieval

To evaluate whether the extended vocabularies could lead to more to greater performance in information retrieval tasks, we compared the number of MIMIC-III annotations and MEDLINE results returned for all non-obsolete subclasses of *Abnormality of the cardiovascular system* (HP:0011025). HP asserts 2,205 labels and synonyms for these classes, while the expanded set of labels numbers 5,336. The results are summarised in Table 3.

**Table 3.** Amount of labels for *Abnormality of the cardiovascular system* (HP:0011025) before and after synonym expansion, and results of the two text mining tasks using them as vocabularies. MEDLINE results are the sum of the number of results returned by each query.

| Vocabulary | Labels | MIMIC-III Annotations | MEDLINE Results |
| --- | --- | --- | --- |
| HP Labels | 2,205 | 1,104 | 8,191,564 |
| Expanded HP Labels | 5,336 | 1,447 | 13,513,342 |

### Patient Characterisation

To evaluate the value of the DO expansion, and to identify whether the additional annotations were informative for the purposes of characterising entities described by biomedical text, we investigated its application within the MIMIC-III dataset [28]. We annotated a sample of 1,000 patient visits using classes from the Disease Ontology (DO) that contained cross-references to ICD-9, both before and after label expansion using the presented algorithm. We then used those annotations to calculate a measure of semantic similarity between the patient visits, and evaluated the rankings with respect to the ICD-9 codes they were annotated with. We used the mean reciprocal rank and the mean average precision to measure how well rankings predicted matching first diagnoses. The results of the ranking task are shown in Table 4, with the expanded vocabulary leading to an increased performance in both cases. To determine whether the result was significantly different, we used the Wilcoxon rank-sum test to compare the ranks of patients with matching first diagnoses, yielding a p-value of 0.0007063.

**Table 4.**
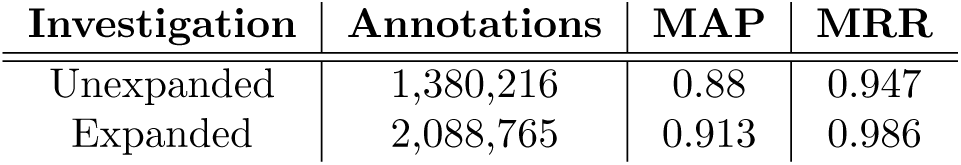
Comparison of the annotations of texts for 1,000 randomly sampled MIMIC-III patient visits before and after expansion, and their associated performance with respect to how predictive semantic similarity scores calculated from the annotations were of shared first diagnosis.

| Investigation | Annotations | MAP | MRR |
| --- | --- | --- | --- |
| Unexpanded | 1,380,216 | 0.88 | 0.947 |
| Expanded | 2,088,765 | 0.913 | 0.986 |

## Discussion

The results clearly demonstrate that for two biomedical ontologies, our approach vastly increases the amount of labels and synonyms available for their terms. Using hypertension as an example, we demonstrated that a range of different ontologies contribute additional synonyms, leading to 25 new unique labels for the term. By leveraging these we can effectively enrich vocabularies for terms.

While we only manually validated a small subset of terms from HP, this indicated a fairly high precision for candidate terms. Through analysis of the false positives, we found that many of them were caused by errors in the ontologies that the synonyms were sourced from. For example, several synonyms for *motor aphasia* (HP:0002427) were marked as incorrect since they refer to dysphasia, including “Broca Dysphasia.” Aphasia and dysphasia are different conditions. The first refers to a partial loss of language, and the latter to a full loss of language. All of these incorrect synonyms were sourced from *Aphasia, Broca* (MESH:D001039) in MESH.

Though this is not reflected in the results, we also found during the development of the algorithm that certain ontologies produced consistently incorrect synonyms. Several of these ontologies are meta-ontologies, automatically constructed from several ontologies using alignment and integration methods, and it’s possible that errors in that process were the cause of the incorrect synonyms. Certain annotation properties were also incorrectly detailed by the AberOWL API as being labels, such as *europe pmc* and *kegg compound*. Candidate synonyms defined by problematic ontologies or matching the list of annotation properties are automatically removed. Expansion of the list of ontologies discluded from the sources for labels might further improve the precision of the algorithm, but may potentially come at the cost of correct synonyms.

Furthermore, the manual validation revealed that many of the returned synonyms were in non-English languages. While OWL ontologies do allow for parameters that distinguish which language the property is in, AberOWL does not index them. Therefore, it is not currently possible to distinguish between English and non-English synonyms. These items were marked as ambiguous, and not counted in the overall precision. This could also be controlled partially by discluding additional ontologies from results. For example, WHOFRE is a non-ontology mapping of French vocabulary to UMLS.

For any uses, where a reduced vocabulary accuracy is not acceptable, the algorithm should be used as a candidate label generator, to be checked by a domain expert before further use.

We also demonstrated that that our expansion of the HP vocabulary increases the amount of data returned by two information retrieval tasks, for a subset of cardiovascular terms. We have not, however, shown in the information retrieval experiment whether the extra information returned is useful or relevant. We can assume, based on our manual validation results that some of the additional data returned are incorrect, though most should be correct.

Finally, our experimental validation showed a clear and significant increase for a patient stratification task over MIMIC-III. This indicates that for certain tasks, our approach can increase the quality of entity characterisations gained by information extraction, and in turn the power of ontology-based analyses, even without manual validation of the produced labels.

### Limitations and Future Work

The most important potential limitation of the algorithm itself is that it violates the notion that the IRI of a concept uniquely identifies it, rather than its name. This is due to the fact that OWL ontologies do not follow the unique name assumption. False positives, in theory, could be generated by a lexical match on a homonym, which then has different synonyms itself. We believe, however, that this effect should be limited in the case of a highly specific biomedical language. Furthermore, any such error would be most likely be mitigated by the dataset context limitation. For example, synonyms derived from different contexts, incorrectly associated with a medical concept, are unlikely to be present within clinical letters. We further limit this effect by only using the first label, reasoning that the most unique synonym will be included as a synonym.

False synonyms could also be removed on the basis of a corpus search. For example, if a candidate synonym never, or, at least, rarely, appears in the same document as another label, used for this term across a literature corpus, it’s possible that it refers to a different concept from a disjoint context. This could also be performed by analysing the metadata of text corpora. For example, if two terms are never, or, at least, rarely, associated with literature from the same journals, the same field, or the same content tags, it’s possible they have different meanings. In a further study, we would investigate whether synonymy can be identified using word embeddings.

While equivalency returns fewer synonyms, and not necessarily many that are unique from the ones gained by lexical matching, they can also be treated with a higher level of confidence. For this reason, using only this method could be considered as a parameter in the case that a higher accuracy is required.

## Conclusions

We have demonstrated that an inter-ontology approach to vocabulary expansion is a powerful method for adding informative labels and synonyms to terms used in text mining. These synonyms are found with a fairly high precision, and lead to a greater rate of document retrieval in clinical and literature settings. Most importantly, we have shown that the approach improves the power of an ontology-based characterisation and analysis of patients via clinical text.

## Methods

All files described in the validation (excluding the MIMIC-III data files), along with the commands necessary to repeat the experiments are available at https://github.com/reality/synonym_expansion_validation/.

### Algorithm

We implemented the algorithm as a module in the Komenti semantic text mining framework using the Groovy programming language [29]. It makes use of the AberOWL API [19] for label matching and semantic queries, documented at http://www.aber-owl.net/docs/.

OWL ontologies use a number of conventional annotation properties to define labels and synonyms. These span a range of confidence and degree of synonymy. In this paper, we consider frequently used annotation properties, summarised in Table 5. These are the annotation properties consolidated into the ‘synonym’ property by the AberOWL API. Another oboInOwl synonym, *hasRelatedSynonym* is excluded, because the labels provided by these synonyms are too imprecise.

**Table 5.** Summary of conventionally used annotation properties considered in this experiment. Definitions come from the description of the annotation properties in their respective top-level ontologies.

| Annotation Property | Identifier | Definition |
| --- | --- | --- |
| label | rdfs:label | “a human-readable version of a resource’s name [30].” |
| altLabel | skos:core#altLabel | “An alternative lexical label for a resource [31].” |
| has_exact_synonym | hasExactSynonym | “An alias in which the alias exhibits true synonymy [32].” |
| has_narrow_synonym | hasNarrowSynonym | “An alias in which the alias is narrower than the primary class name. Example: pyrimidine-dimer repair by photolyase is a narrow synonym of photoreactive repair [32].” |
| has_broad_synonym | hasBroadSynonym | “An alias in which the alias is broader than the primary class name. Example: cell division is a broad synonym of cytokinesis [32].” |
| alternative term | IAO_0000118 | “An alternative name for a class or property which means the same thing as the preferred name (semantically equivalent) [33].” |

### Manual Validation

To evaluate the performance of the algorithm, we randomly selected 500 classes from the expanded version of HP for manual validation. Synonyms already asserted by HP were removed from the set, because they were already assumed to be correct, and would not contribute to measuring the performance of the synonym expansion algorithm. A clinical expert (WB) marked each synonym as correct, incorrect, or ambiguous. The expert was asked to answer correctly or incorrectly on the basis: “if a patient has *synonym*, would it also be true that they have *original label* ?” Entries were marked as ambiguous if the synonym was in a different language, or the validator otherwise did not have enough knowledge of the phenotype to determine whether or not the synonym was correct.

### Information Retrieval

We used the Komenti semantic text mining framework, which implements Stanford CoreNLP’s RegexNER [34] to annotate 1,000 randomly sampled entries from the NOTEEVENTS table in MIMIC-III (MIMIC) [35]. MIMIC is a freely available healthcare database, containing a variety of structured and unstructured information concerning around 60,000 admissions to critical care services [28]. We annotated the sample with all subclasses of *Abnormality of the cardiovascular system* (HP:0011025), comparing the number of annotations before and after synonym expansion. This investigation was performed on 17/01/2020.

Using the same set of subclasses of *Abnormality of the cardiovascular system* (HP:0011025), we compared the sum of article counts returned for a disjunctive query of all labels and synonyms for each term, before and after synonym expansion. MEDLINE is a searchable database of literature metadata in the life sciences, containing more than 25 million article references [36]. MEDLINE was queried on the 27/01/2020.

### Patient Characterisation

We sampled 1,000 patient visits from the MIMIC-III (distinct from those used in the information retrieval experiment). We then concatenated all text records for each patient visit from the NOTEEVENTS table into one text file, and pre-processed the text to remove newlines, improve sentence delineation, and lemmatise words. We also retained the primary diagnosis, which was the first listed ICD-9 code in the DIAGNOSES ICD table. These codes are produced by clinical coding specialists, by examining the texts associated with the visit.

We limited the classes considered for our annotation vocabulary to those which DO contained a database cross-reference to ICD-9, of which there were 2,118. This was to reduce noise from terms not represented in ICD-9. We obtained the unexpanded and expanded synonyms for these terms on 08/07/2020. Both sets of labels were also lemmatised (both lemmatised and unlemmatised forms were used for annotation).

The Komenti semantic text-mining framework was used to annotate the text associated with each patient visit. As before, this made use of the CoreNLP RegexNER annotator [34]. Negated annotations were excluded using the komenti-negation algorithm [37]. We then used the set of terms associated with it to produce a semantic similarity matrix for patient visits, using the Resnik measure of pairwise similarity for each annotated term [38], normalised into a groupwise measure using the best match average method [39]. Information content was calculated using the probability of the term appearing as an annotation in the totality of the set of annotations [38]. The similarity matrix was computed using the Semantic Measures Library [40].

We evaluated the similarity matrix using mean reciprocal rank and mean average precision to measure performance in predicting shared primary patient diagnosis. A true case was considered to be whether a pair of patient visits had the same primary diagnosis (as per the MIMIC-III database). For mean average precision, we considered only the 10 most similar patients for each patient. The p-value was calculated using the built-in *wilcoxon.test* function of R version 3.4.4 [41].

## Author’s contributions

LTS conceived of the study, performed the experiments, implemented the software, and wrote the first draft the manuscript. RH conceived of the patient characterisation experiment. WB performed data validation and contributed to evaluation of results. RH and GVG contributed to the manuscript. GVG, RH, and SB supervised the project. All authors revised and approved the manuscript for submission.

## Acknowledgements

The authors would like to acknowledge Dr Andreas Karwath for advice on evaluating ranking algorithms. We would further like to thank Dr Paul Schofield and Dr Egon Willighagen for advice concerning an earlier version of the experiment, particularly surrounding precision and error. GVG and LTS acknowledge support from support from the NIHR Birmingham ECMC, the NIHR Birmingham SRMRC, Nanocommons H2020-EU (731032), OpenRisknet H2020-EINFRA (731075) and the NIHR Birmingham Biomedical Research Centre and the MRC HDR UK (HDRUK/CFC/01), an initiative funded by UK Research and Innovation, Department of Health and Social Care (England) and the devolved administrations, and leading medical research charities. The views expressed in this publication are those of the authors and not necessarily those of the NHS, the National Institute for Health Research, the Medical Research Council or the Department of Health.

